# Heterogeneity and excitability of *BRAF*^*V600E*^-induced tumors is determined by PI3K/mTOR-signaling state and *Trp53*-loss

**DOI:** 10.1101/2021.02.22.432030

**Authors:** Silvia Cases-Cunillera, Karen M. J. van Loo, Julika Pitsch, Anne Quatraccioni, Sugirthan Sivalingam, Paolo Salomoni, Dirk Dietrich, Susanne Schoch, Albert J. Becker

## Abstract

**Background:** Developmental brain tumors harboring *BRAF*^*V600E*^ somatic mutation are diverse. Here, we describe molecular factors that determine *BRAF*^*V600E*^-induced tumor biology and function.

**Methods:** Intraventricular *in utero* electroporation in combination with the piggyBac transposon system is employed as a tool to generate developmental brain neoplasms. *In vivo* tumor growth is monitored by using the infrared fluorescent protein (iRFP). Lineage inference is carried out by using the Brainbow transgene. Neural activity from tumor slices is assessed by multielectrode array. RNA sequencing is exploited to analyze the induced neoplasms at the transcriptomic level.

**Results:** *BRAF*^*V600E*^ in murine neural progenitors only in concert with active PI3K/mTOR-signaling through constitutively phosphorylated Akt-kinase (*p*Akt) elicits benign neoplasms composed of enlarged dysmorphic neurons and neoplastic astroglia recapitulating ganglioglioma (GG). Purely glial tumors partially resembling polymorphous low-grade neuroepithelial tumors of the young (PLNTYs) emerge from *BRAF*^*V600E*^ alone. Additional somatic *Trp53*-loss is sufficient to induce anaplastic GGs (aGGs) with glioneuronal clonality. Functionally, only *BRAF*^*V600E*^/*p*Akt tumors intrinsically generate substantial neuronal activity and show enhanced relay to adjacent tissue conferring high epilepsy propensity. In contrast, PLNTY- and aGG-models lack significant spike activity, which appears in line with the glial differentiation of the former and a dysfunctional tissue structure combined with reduced neuronal transcript signatures in the latter.

**Conclusion:** mTOR-signaling and *Trp53*-loss critically determine the biological diversity and electrical activity of *BRAF*^*V600E*^-induced tumors.

**Key points:**

- IUE of BRAF^V600E^ and activation of mTOR leads to ganglioglioma (GG)-like tumors, while BRAF^V600E^ alone give rise to PLNTY-like neoplasms.
- Anaplastic GGs depend on the Trp53 deletion in combination to BRAF^V600E^ and PI3K-mTOR signaling cascade.

**Importance of the Study:** Glioneuronal tumors are challenging with respect to biological behavior and seizure emergence. While *BRAF*^*V600E*^ in murine neural precursors induces oligoid tumors, it requires an overactivation of PI3K/mTOR-signaling for the development of hyperexcitable gangliogliomas and additional *Trp53*-loss for anaplastic transformation.

## Introduction

Activating *BRAF*^*V600E*^ somatic mutation has been detected in a large spectrum of developmental low-grade glial and glioneuronal brain tumors ^1–5^. *BRAF*^*V600E*^ is particularly frequent in up to approximately 60% of gangliogliomas (GGs) and has been observed in both, dysmorphic neuronal and neoplastic astroglial tumor cells ^6^. GGs are typical pediatric neoplasms and represent the most frequent tumor entity in patients undergoing epilepsy-surgery ^7,8^. Recent data demonstrated that intraventricular *in utero* electroporation (IUE) of the murine *BRAF*^*V600E*^ equivalent (*Braf*^*V637E*^) during early brain development was sufficient to induce cytoarchitectural abnormalities of mutant neurons, increased numbers of astro- and oligodendroglia and seizures in mutant mice ^9^.

However, *BRAF*^*V600E*^ is present in a large variety of purely glial including polymorphous low-grade neuroepithelial tumor of the young (PLNTY) ^5^ as well as glioneuronal tumors with malignant features and differences in seizure propensity. The molecular factors conferring this heterogeneity are unresolved. Recent reports have suggested that the PI3K/mTOR pathway is activated in human *BRAF*^*V600E*^-positive GGs ^10–13^. Despite the fact that most GGs behave biologically benign, anaplastic variants (aGGs) occur and represent onco- and epileptological challenges. The significance of *Trp53*-mutations has remained controversial in the context of human aGGs ^14–16^. A recent study reported *Trp53*-mutations as relevant prognostic parameter for seizure control in human aGGs ^17^.

Here, we have scrutinized in IUE-induced somatic *BRAF*^*V600E*^-positive mouse developmental brain tumors, whether active PI3K/mTOR-signaling mediated through constitutively phosphorylated Akt-kinase (*p*Akt) as well as *Trp53*-loss confer heterogeneity and distinct electrical activity to emerging tumors.

## Methods

### Experimental model and subject details

#### Patient samples and experimental animals

Human *BRAF*^*V600E*^-positive tumors (*n* = 3 GGs and PLNTY, each) were from resections within the University of Bonn Medical Center Neurosurgery Program. Written consent was obtained from all patients with respect to the use of brain tissue for additional studies. All procedures were conducted in accordance with the Declaration of Helsinki and approved by the Ethics Committee of the University of Bonn Medical Center. All experiments involving animals were performed in accordance with the guidelines of the European Union and the University of Bonn Medical Center Animal Care Committee.

#### Intraventricular *In utero* electroporation

IUE was performed as described previously ^18^. The DNA solution (at a final concentration of 1.5 μg/μl) was injected into the lateral ventricle of mouse embryos with microcapillaries and expelled by pressure with a microinjector (Picospritzer III, General Valve Corporation). The electrodes were placed to target cortical progenitor cells and five electric pulses of 45 V were delivered using the CUY21 SC Square Wave Electroporator (Nepa Gene).

#### Neuropathology in human and mouse brain tissue

Mouse brain tissue was fixed in 4% formaldehyde (Sigma Aldrich) overnight at 4 °C and embedded in paraffin. Formalin-fixed paraffin embedded (FFPE) brain tissue blocks were sectioned at 4 μm with a Microtome (HM-335-E, GMI) and dried overnight at 37 °C. See **Supplementary Materials and Methods** for the histological and immunohistochemical analysis of FFPE tissue.

#### RNA sequencing and bioinformatic analysis

After RNA isolation, library preparation was performed using QuantSeq 3′ mRNA-Seq Library Prep kit (Lexogen). Samples were subjected to sequencing on a HiSeq 2500 sequencer (Illumina) with 1×50 bp single-end reads and a coverage of 10.000 reads per sample (see **Supplementary Materials and Mehods**). All read counts datasets that support the data of this study are available upon request. A FDR threshold of 5% was chosen to identify differentially expressed (DE) genes.

#### Near-infrared *in-vivo* imaging

Mice IU-electroporated with CAG-iRFP^713^ and the respective oncogenes were anesthetized and the skull was exposed. Pearl Impulse Small Animal Imaging System (Li-COR Biosciences GmbH) was used to perform near-infrared imaging (at P10, P20 and P45) and the fluorescent signals were quantified as described previously ^19^.

#### MEA recordings and data analysis

Acute brain slices were transferred on a multielectrode array (MEA) plate (see **Supplementary Materials and Methods**). The slices were kept onto the electrodes using a platinum grid and were incubated in aCSF oxygenated with carbogen for the whole recording. The spontaneous activity of the slices was recorded with the AxIS software (Axion Integrated Studion Navigator 1.5., Axion Biosystems) for 15 min.

#### Statistical analysis

Statistical analysis and graphs were performed by using GraphPrism, Igor 64 and R Studio. All results are plotted as ± SEM. For each IHC Ki67 and phospho-S6 ribosomal protein (pS6) staining, the labeling index (LI) of 10 tissue sections was quantified from the mean values. Within all models, the LI was statistically compared between different immunostainings using One-way ANOVA, Bonferroni′s Multiple Comparison Test ^11^.

#### Resource availability

Required data and further information should be directly requested and will be provided by the Lead contact, Albert J. Becker.

## Results

### Architecture of *BRAF*^*V600E*^-induced brain tumors is determined by PI3K/mTOR pathway activation

The spectrum of *BRAF*^*V600E*^-positive tumors comprises glioneuronal/GG and glial/PLNTY entities (**Fig. 1A,** left panel). We observed the phospho-ribosomal protein S6 (*p*S6) as downstream effector of PI3K/mTOR pathway signaling present in GGs (**Fig. 1A**, right panel upper part) ^13^ but virtually absent in PLNTYs (**Fig. 1A**, right panel lower part, *n* = 3 each). It is so far unresolved, whether PI3K/mTOR pathway activity with its surrogate *p*S6 is relevant for the emergence of glioneuronal lesions or e.g. a consequence of frequently associated epileptiform activity ^13,20^. To address a potential developmental role of PI3K/mTOR signaling in *BRAF*^*V600E*^-induced brain tumors, we used IUE with the piggyBac transposon system at E14 (**Supplementary Fig. S1A**) ^21^. IUE of the piggybac transposase (pBase) together with *BRAF*^*V600E*^ and/or constitutive *p*Akt piggyBac transgenes results in selective integration of transposons carrying *BRAF*^*V600E*^*/p*Akt in IU-electroporated cells ^22,23^ and subsequently transference to successive progenies of the initially IU-electroporated precursor cells.

**Figure 1.**
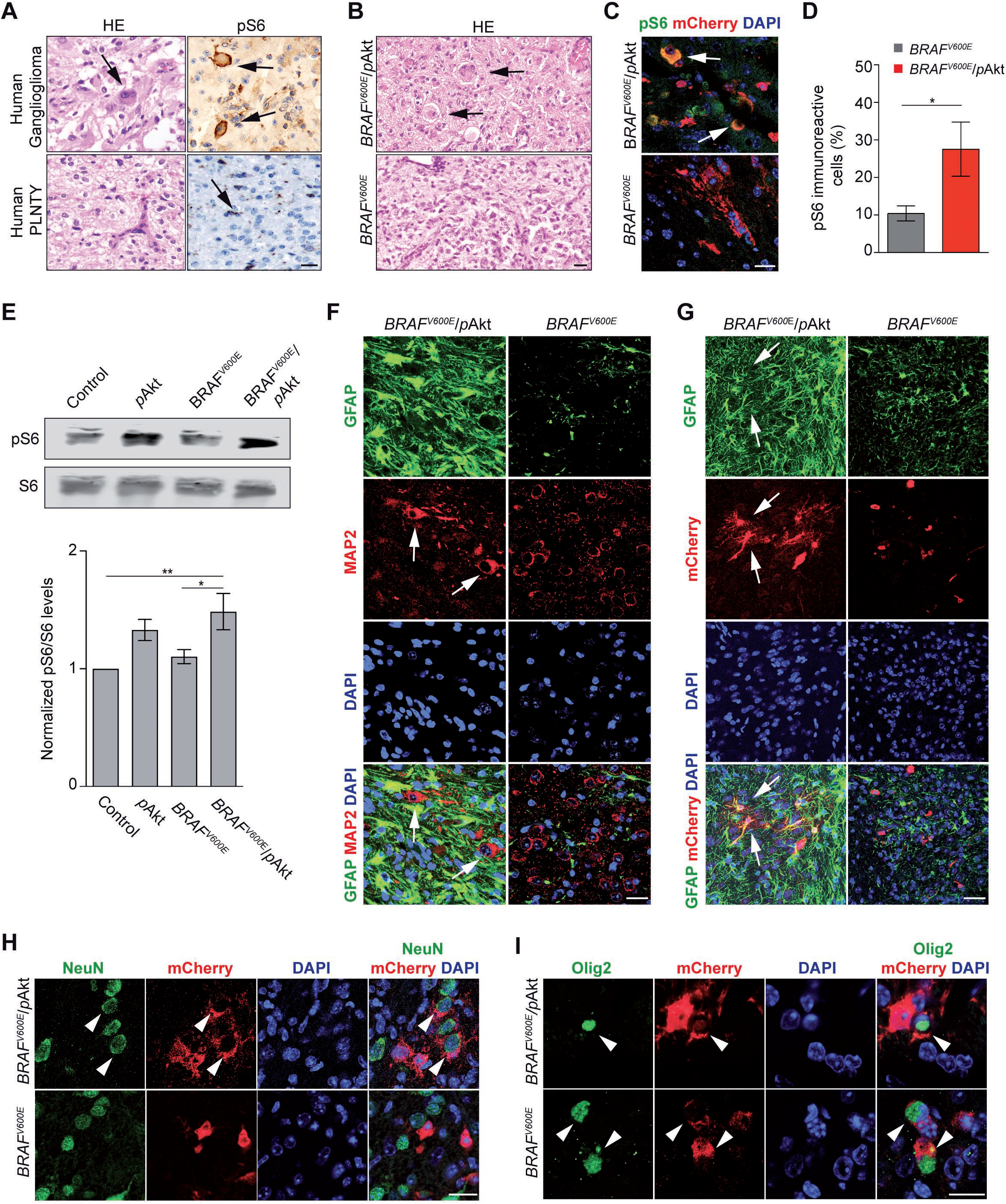
Level of mTOR pathway activation fundamentally impacts *BRAF*^*V600E*^-induced tumors characteristics. **(A)** Representative human GG (upper panels) versus monomorphic oligoid PLNTY (lower panels). Scale bar, 100 μm. (**B)** HE-stained sections of *BRAF*^*V600E*^(1-hit)- and *BRAF*^*V600E*^/*p*Akt(2-hit)-IUE brains (P110). **(C)** Expression of *p*S6 in *BRAF*^*V600E*^ and *BRAF*^*V600E*^/*p*Akt-induced tumors (*n* = 5). Scale bar, 25 μm. **(D)** Quantification of *p*S6-immunoreactive cells in the 2-versus 1-hit tumors (*n* = 5). Two-tailed unpaired t-test, *p < 0.05). **(E)** Normalized ratio between pS6/S6 protein levels of NS20Y cells transfected with CAG-mCherry (Control), CAG-*p*Akt-mCherry (*p*Akt), CAG-*BRAF*^*V600E*^-mCherry (*BRAF*^*V600E*^) and both (*BRAF*^*V600E*^/*p*Akt). One-Way ANOVA followed by Tukey′s multiple comparison test, *p < 0.05; **p < 0.01, (*n* = 4). **(F)** *BRAF*^*V600E*^/*p*Akt-induced tumors reveal an admixture of large dysplastic MAP2-positive neurons (white arrows) and a dense GFAP-positive astroglial matrix. 1-hit model tumors composed by small isomorphic MAP2-positive elements. **(G)** Immunofluorescent labeling of GFAP by a fraction of mCherry-positive cells in *BRAF*^*V600E*^/pAkt tumors (white arrows) indicates the presence of astroglial tumor cells in concert with GFAP-positive/mCherry-negative reactive astroglia. In 1-hit tumors, GFAP-positive tumor cells are absent. **(H)** NeuN- and mCherry (*BRAF*^*V600E*^/*p*Akt)-positive large neurons in the 2-hit model (white arrowheads) versus small and circular shaped mCherry-positive but NeuN-negative cells in the 1-hit model. **(I)**Co-expression of Olig2 and mCherry in both, 1- and 2-hit model tumors (white arrowheads). Scale bars, 25μm.

Intriguingly, only co-IUE of *p*Akt and *BRAF*^*V600E*^ (further designated as *BRAF*^*V600E*^/*p*Akt or 2-hit model; **Supplementary Fig. S1E**) results in tumors harboring enlarged, sometimes binucleated dysmorphic neurons, intermingled with process-rich astroglial cells and strong *p*S6 expression (*n* = 5, **Fig. 1B** and **1C**, upper panels; **Fig. 1D**). All animals IU-electroporated with *BRAF*^*V600E*^ (1-hit model; **Supplementary Fig. S1D**) developed circumscribed lesions composed of oligoid cells and lacked significant expression of *p*S6 (*n* = 8**; Fig. 1B** and **1C,**lower panels; **Fig. 1D**), whereas none of the control animals (*n* = 8; **Supplementary Fig. S1B**) showed any tumor. IUE of only *p*Akt (**Supplementary Fig. S1C**) at E14 did not result in detectable tumors until P110 (data not shown; *n* = 6). Complementary immunoblot of neuroblastoma NS20Y cells revealed that only the combination *BRAF*^*V600E*^/*p*Akt leads to increased pS6 levels compared to basal and *BRAF*^*V600E*^-stimulated conditions (**Fig. 1E**).

Subsequent immunofluorescence analysis of 2-hit tumors revealed a *glioneuronal* phenotype composed of enlarged dysmorphic neuronal shaped cells positive for the microtubule-associated protein 2 (MAP2) interspersed in a dense glial fibrillary acidic protein (GFAP)-positive fibrillary matrix (**Fig. 1F**, left panels). In contrast, 1-hit neoplasms contained MAP2 positive cells with a monomorphic shape and only few non-neoplastic, thus reactive GFAP-positive astrocytes (**Fig. 1F**, right panels). Additional immunofluorescent analyses demonstrated co-localization between GFAP and mCherry-IU-electroporated cells in 2-hit tumors (**Fig. 1G**, left panels) and we observed mCherry-IU-electroporated cells positive for nuclear Ki-67 expression (**Supplementary Fig. S2**), suggesting a neoplastic astroglial component. Contrarily, the lack of mCherry-IU-electroporated cells positive for GFAP in 1-hit neoplasms identified GFAP-positive cells as reactive astrocytes (**Fig. 1G and Supplementary Fig. S2**, right panels). *BRAF*^*V600E*^/*p*Akt-mCherry, but not *BRAF*^*V600E*^-targeted cells, co-localize with the neuron-specific nuclear protein NeuN, indicating their neuronal phenotype (**Fig. 1H**). Both, 1- and 2-hit tumors reveal mCherry- and oligodendrocyte transcription factor 2 (Olig2)-positive tumor cells (**Fig. 1I**). Olig2 expressing oligoid-shaped cells recapitulate key features of human PLNTY ^5^. These results indicate that *BRAF*^*V600E*^ alone in E14 neural progenitors elicits tumors with oligodendroglial, thus PLNTY-like features, as well as in concert with robust PI3K/mTOR-pathway signaling glioneuronal neoplasms recapitulating GG features. *p*Akt-induced PI3K/mTOR-pathway signaling acts as non-neoplastic but phenotypically modifying factor of *BRAF*^*V600E*^-induced tumors. Of note, 1- and 2-hit tumors lacked increased mitoses and thus, appeared benign. In GGs, anaplasia of the astroglial component is an enigmatic issue ^24^.

### Anaplasia of *BRAF*^*V600E*^/*p*Akt-induced tumors through *Trp53* loss

The significance of anaplastic histological features in GGs for patient survival has remained controversial ^25–27^. Loss-of-function *Trp53* mutations have been detected in human aGGs but the pathogenetic impact has remained unclear ^15,28^. To study this aspect, we have modified the 2-hit model by co-IU-electroporating *BRAF*^*V600E*^, *p*Akt and Cre at E14 in *Trp53*^*loxP/loxP*^ mice (further designated as *BRAF*^*V600E*^/*p*Akt/*Trp53*^*KO*^- / 3-hit-model; **Supplementary Fig. S1F**). *BRAF*^*V600E*^/*p*Akt/*Trp53*^*KO*^ introduced into neural progenitors resulted in diffusely infiltrating tumors with high cellularity and Ki67-labeling index of the astroglial, thus neoplastic malignant component (*n* = 19; **Fig. 2A and 2B**). The tumors retained GG-features by the presence of large, dysmorphic neurons (**Fig. 2B and 2C**). The astroglial tumor component appeared malignant due to high pleomorphism, density and dynamic proliferation (**Fig. 2D,** *arrows*). Of note, IUE of only Cre at E14 in *Trp53*^*loxP/loxP*^ mice did not result in any emerging tumor (*n* = 5; data not shown). Thus *Trp53*-loss does not elicit aGG-like tumors independently from *BRAF*^*V600E*^/*p*Akt, but in the present context acts as modifier that leads to the acquisition of anaplastic features of the emerging neoplasms.

**Figure 2.**
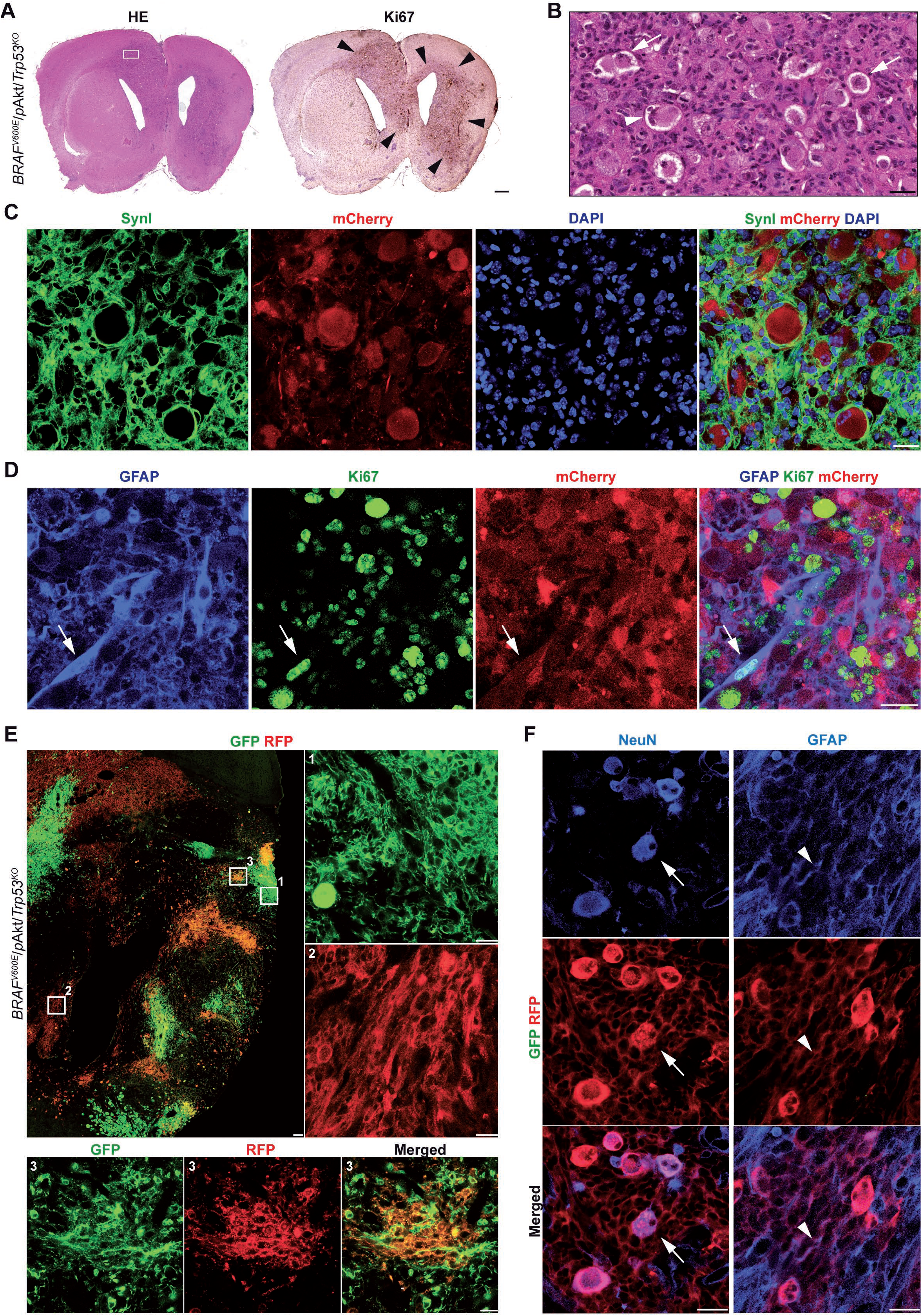
Anaplastic phenotype and clonal architecture through *Trp53*-loss in developmental *BRAF*^*V600E*^/*p*Akt-induced tumors. **(A)** Coronal view of a *BRAF*^*V600E*^/*p*Akt/*Trp53*^*KO*^ mouse brain (P40) harboring a large hemispheric tumor with prominent mass effect (HE-staining). Note the tumor cells positive for the proliferation-related antigen Ki67 (arrow heads); scale bar, 500μm. **(B)** Enlarged area of the tumor shown in **A** (square) demonstrates the presence of dysmorphic (arrows), occasionally binucleated (arrow head) neurons in an astroglial matrix with high cellularity and pleomorphism, thus reflecting anaplastic features. **(C)** Immunofluorescence confirms the SynI-positive neurons as mCherry-positive tumor components. **(D)** The GFAP-/mCherry-positive astroglial tumor cell component (white arrows) reveals a high fraction of Ki67-expressing elements as sign of anaplasia. **(E)**Representative section of a *BRAF*^*V600E*^/*p*Akt/*Trp53*^*KO*^/*Brainbow* tumor at P20 (*n* = 5 in total); scale bar, 100 μm. Right and low panels show high-magnification images of white squares 1, 2 and 3 from the overview image. **(F)** Neurons (arrows; NeuN-IHC) and astroglial (arrow heads; GFAP-IHC) cells in a brainbow RFP-labeled tumor clone, suggesting an offspring of both cell types from a pluripotent neural precursor; scale bars, 25 μm.

### ‘Glioneuronal clonality’ of *BRAF*^*V600E*^/*p*Akt/*Trp53*^*KO*^ tumor cell populations

Particularly for aGGs, the glioneuronal character has been questioned due to the fact that the glial tumor fraction recapitulates malignant features resembling glioblastoma multiforme such that dysmorphic neurons and neoplastic astroglia might derive from different rather than identical precursor cells. To address this issue, we used a multicolor system of donor plasmids driving expression of enhanced green fluorescent protein (eGFP), mOrange and mKate2 under control of the CAG promoter creating multicolor clonal labeling (Brainbow 3.0 allele cloned into the piggyBac plasmid; **Supplementary Fig. S3A**) ^29,30^. We next IU-electroporated the piggyBac-Brainbow construct when eliciting 3-hit model tumors (*n* = 5; further designated as *BRAF*^*V600E*^/*p*Akt/*Trp53^KO^*/*Brainbow*; **Supplementary Fig. S3D**). Multicolor labeling of tumor cells at P20 allowed the visualization of tumor cell clonality (**Fig. 2E**). The intermingled distribution of differently colored astroglia indicates parallel clonal expansion and mixing of tumor cells (**Fig. 2E**). Overall, distinct colors were always labeling neurons (representative images **Fig. 2F;** left panels) *and* astroglia (representative images **Fig. 2F;** right panels), indicating that the present IUE generally hits *glioneuronal* progenitor cells enforced by the genetic manipulation to give rise to both, dysmorphic neuronal and glial tumor cell fractions ^31^ and argues against an ontological concept of GGs based tumorigenesis by neoplastic transformation of astroglia within a dysplastic precursor lesion. Thus, dysmorphic neurons in concert with malignant astroglia in 3-hit tumors do not only represent co-incidental phenomena but rather shape an integrated neoplastic unity.

### Growth dynamics and survival kinetics reflect tumor genetics

Concerning biological behavior, both 1- and 2-hit models revealed favorable survival kinetics (*n* = 8, with 87.5% of the mice still being alive at the age of P110 for *BRAF*^*V600E*^-positive tumors, 88.89% survival at P110 (*n* = 9) for *BRAF*^*V600E*^/*p*Akt-positive tumors; **Fig. 3A**) and very low Ki67-labeling proliferation indices (**Fig. 3B** and **3C**). In contrast, anaplastic histological features of 3-hit tumors were reflected by poor survival up to only P70 (*n* = 19; **Fig. 3A**) and an extensively increased Ki67-immunoreactive index (**Fig. 3B** and **3C**).

**Figure 3.**
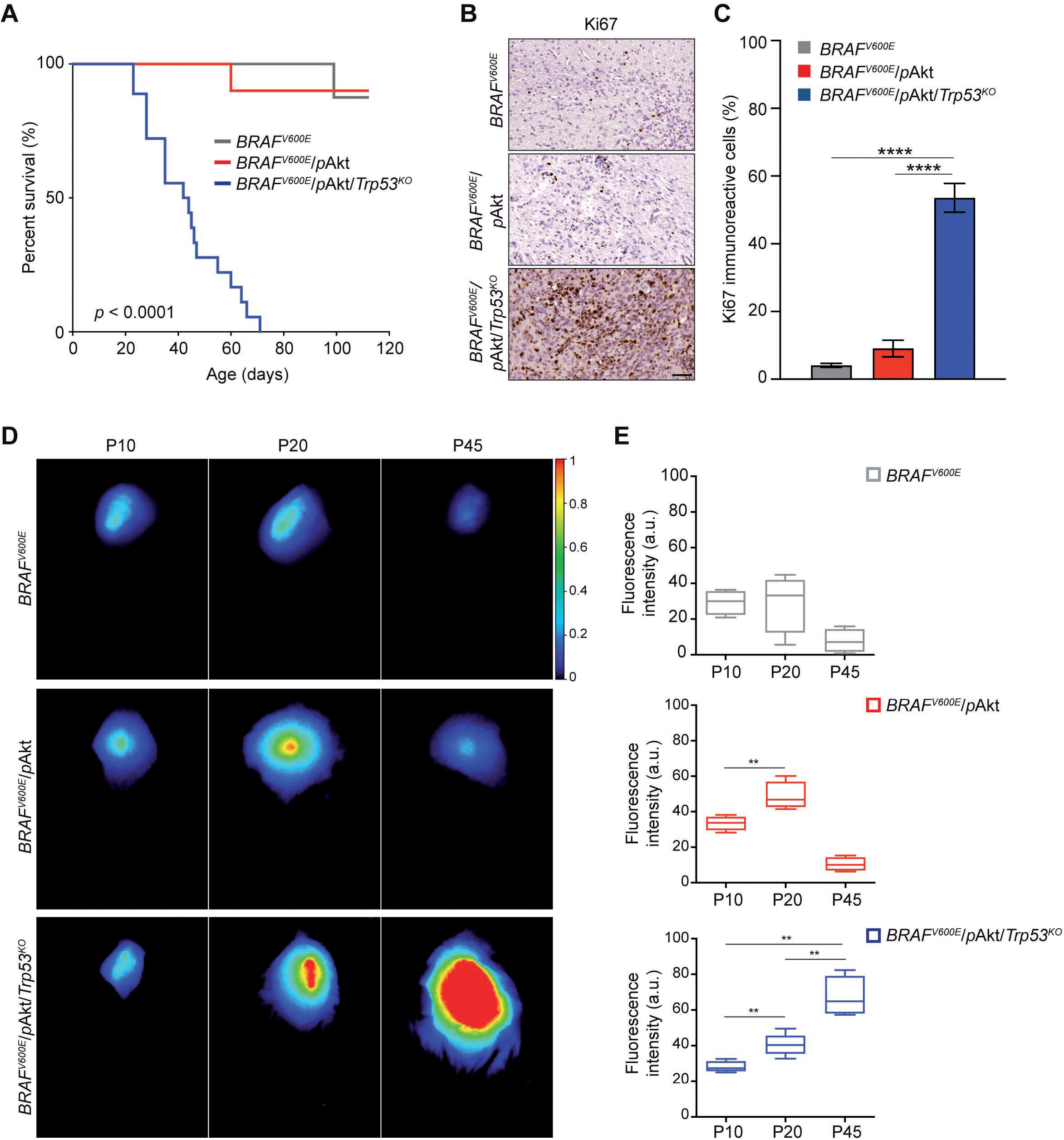
Distinct *in-vivo* growth kinetics of GG-models. **(A)** Kaplan-Meier survival curves of mice IU-electroporated at E14 with *BRAF*^*V600E*^ (*grey* line; *n* = 8), *BRAF*^*V600E*^/*p*Akt (*red* line; *n* = 9) and *BRAF^V600E^/p*Akt*/Trp53^KO^* (*blue* line; *n* = 19); Log-rank test, p < 0.0001. **(B)** Representative photomicrograph of Ki67-stained brain sections for the different tumor models. **(C)** Percentage of proliferating, neoplastic cells within the tumor area. Note significantly increased proliferation activity in 3-hit tumors compared to both other models (*N* = 3 – 5; *n* = 10). **(D)** Representative *in-vivo* iRFP brain tumor signals of 1-, 2- and 3-hit tumor mice detected at P10, P20, and P45 (color bar - total fluorescence efficiency in pseudo-color). **(E)** Quantification of iRFP signals of the different tumor variants (*n* = 4 – 6). Regions of interest (ROIs) were defined above the tumor region and the fluorescence intensity was defined in arbitrary units (a.u.). One-way ANOVA followed by Tukey′s multiple comparisons test; * p < 0.05; **p < 0.01, ****p < 0.0001.

Despite the fact that differences in survival rate and proliferation of the tumor models indicate distinct biological behavior, we sought to analyze *in-vivo* whether different molecular architectures substantially impact tumor growth kinetics by IUE of a near-infrared fluorescent protein (iRFP) plasmid (CAG-iRFP^713^) ^32^. *In-vivo* near-infrared imaging at P10, P20 and P45 allowed to follow growth dynamics in the individual tumor models (**Fig. 3D**). Intriguingly, *BRAF*^*V600E*^-positive neoplasms represent early developmental tumors since the growth dynamics are apparently exhausted already at P10 (**Fig. 3E**). The initial oncogenic stimulus mediated by the mutant BRAF^V600E^ variant may rapidly convert into oncogene-induced interruption of proliferation and differentiation as tumor-related phenomenon that parallels previous observations coined ‘*BRAF*^*V600E*^-induced senescence’ in human neural stem cells and progenitors ^33^. In contrast, 2-hit tumors preserve growth dynamics for a longer time period until P20, whereas in mice harboring 3-hit tumors, dynamic expansion even remains present throughout the entire observational period (**Fig. 3E**).

### Impaired neuronal signaling in concert with invasive glial RNA signatures operative in *BRAF*^*V600E*^/*p*Akt/*Trp53*^*KO*^ tumors

Next, we pursued to understand whether 3-hit tumors would recapitulate transcript features of high grade gliomas or glioneuronal tumors ^9,34,35^ or alternative integrate characteristics of both, thereby creating a unique RNA-fingerprint. RNA-(seq)uencing of *BRAF*^*V600E*^/*p*Akt/*Trp53*^*KO*^ tumor and grey-/white matter matched control (ctrl) tissues (*n* = 4 per group; P40) identified a substantial number of genes to be differentially expressed (3141 genes in- versus 2700 genes decreased in expression in tumor tissue; **Fig. 4A**). Principal component analysis (PCA) of the RNA-seq data clearly separated the 3-hit tumors from control tissue samples (**Fig. 4B**). Intriguingly, GO enrichment analysis revealed that the term with most pronouncedly induced signature in fact relates to inflammation and is accompanied by augmented transcripts with biological functions that reflect malignant glioma characteristics including proliferation, invasion and neovascularization (**Fig. 4C**). Transcripts reduced in tumor tissue grouped under GO-terms related to neuronal homeostasis and signal transduction (**Fig. 4C**). This observation rather reflects true expression regulation rather than simply different tissue composition between tumor and control samples since a significant portion of neuronal transcripts was unchanged or even increased between tumor and control tissue (**Supplementary Table S1)**, We, finally, analyzed differential expression of ion channels and neurotransmitter receptors in detail. Heat map clustering revealed an overall lower expression of distinct families of channels and receptors (**Fig. 4D**). In each family only sets of transcripts showed robustly reduced transcript patterns whereas other subunits were unchanged in tumor tissue (**Fig. 4E-J**). Considering the particular morphological and molecular features of the aGG model, we aimed to understand whether these tumors differ from the low-grade *BRAF*^*V600E*^-induced neoplasms in functional, excitability-related terms.

**Figure 4.**
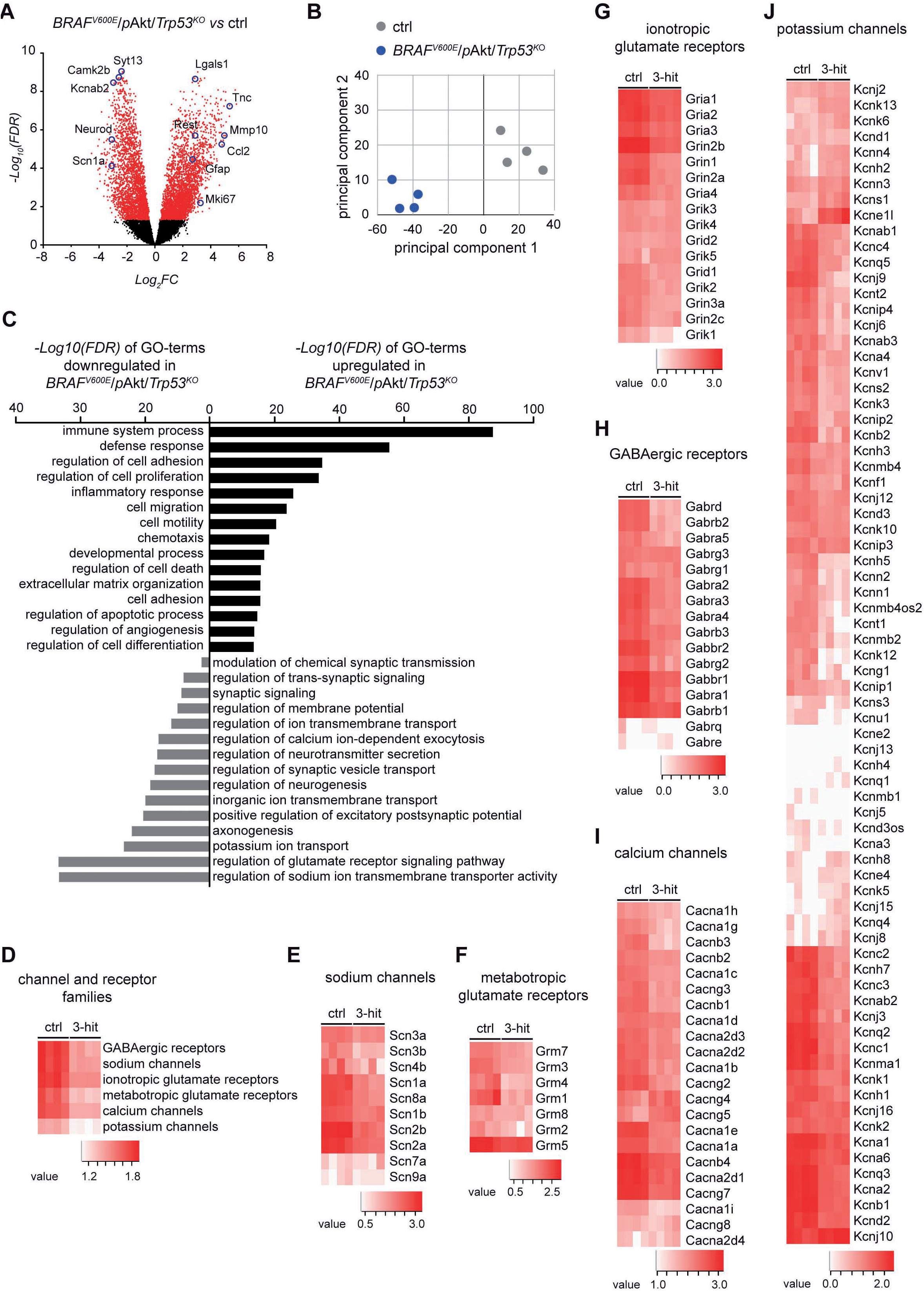
Transcriptomic analysis of *BRAF*^*V600E*^/*p*Akt/*Trp53*^*KO*^-induced tumors reveals malignant glial and non-homeostatic neuronal signatures. **(A)** Volcano plot represents differentially expressed genes in 3-hit tumors compared to grey/white matter matched control (ctrl) tissue (*n* = 4). A total number of 5842 genes were differentially expressed for 3-hit tumors compared to the respective healthy brain counterpart Each gene is colored based on the −log_10_ adjusted *p*-value (FDR). *Red* dots: FDR < 0.05: *black* dots: FDR > 0.05. *Blue* circles indicate a selection of significantly increased or decreased genes, playing an important role in the process of tumorigenesis. **(B)** PCA plot clearly separates 3-hit tumors (*blue* dots) from controls (*grey* dots); transcript signatures of four biological replicates. **(C)** GO-terms based on significantly enriched genes from 3-hit tumors compared to control tissue. **(D-J)** Heatmap visualizations of expression levels of genes coding for synaptic-related channels and receptors.

### Electrical activity patterns differentiate *BRAF*^*V600E*^-tumor variants

*BRAF*^*V600E*^-positive/glioneuronal tumors are particularly epileptogenic and altered neuronal activity in tumor microenvironmental (TME) networks has been recently demonstrated also in malignant gliomas ^9,36–38^. We sought to analyze systematically whether differences in intrinsic excitability as well as TME neuronal networks occur in the different *BRAF*^*V600E*^-positive tumors under study and, therefore, tested electrical activity in acute brain slices containing the tumor at P50-60 using a multielectrode array (MEA) system. Based on the distribution of mCherry-positive cells, electrodes were classified into three categories: i) IUE Tumor core (area with mCherry-positive tumor cells), ii) Peri-IUE (tumor border) and iii) Non-fluorescent tissue (pre-existing brain tissue) (**Fig. 5A** and **5B**). Extracellular activity was recorded by the 64 electrodes of the MEA plate, encompassing the three different categories (**Fig. 5C**). Similar to previous results from low-grade human gliomas ^36^, the lowest frequency of spontaneous activity was observed for all models in the tumor core (**Fig. 5D**). Spontaneous spike activity from slices incubated with artificial cerebrospinal fluid (aCSF) could clearly be resolved within the three different categories and in all groups (**Fig. 5D** and **5E**) and at frequencies comparable to control in the peri-tumoral area and the surrounding tissue (**Fig. 5F**; *yellow* and *blue* bars). However, the groups significantly differed in the number of spikes produced per area in the tumor core region: only *BRAF*^*V600E*^/*p*Akt tumors displayed activity at a level not different to the control whereas both *BRAF*^*V600E*^ only and *BRAF*^*V600E*^/*p*Akt/*Trp53*^*KO*^ appeared almost silent in this central region (*red* bars).

**Figure 5.**
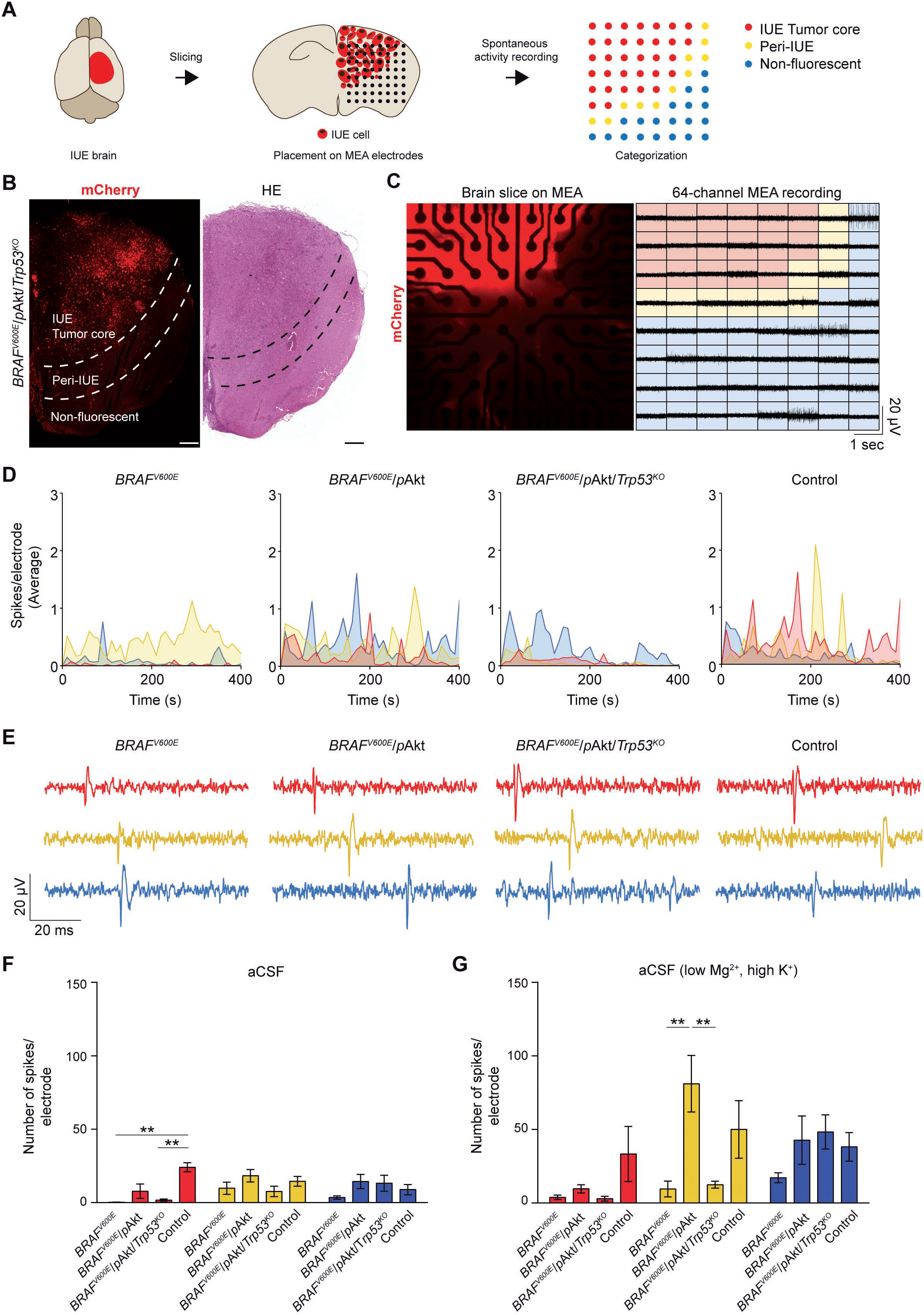
MEA-based recordings of spontaneous neuronal activity of murine GG brain slices. **(A)** Tumor mouse brains were sliced at P50-60 and placed onto a grid of 64 electrodes in a MEA plate. Electrodes were classified according to the area of the slice they contacted: In *red*, electrodes in contact with tissue expressing mCherry (IUE Tumor core); In *yellow*, electrodes in contact to adjacent tissue from areas expressing mCherry (Peri-IUE); In *blue*, electrodes in contact with tissue negative for mCherry fluorescent expression (Non-fluorescent). **(B)** Overview of a coronal slice of a *BRAF*^*V600E*^/*p*Akt/*Tp53*^*KO*^ brain. Scale bar, 500μm. **(C)** *BRAF*^*V600E*^/*p*Akt/*Tp53^KO^* tumor slice on the MEA grid consisting of 64 electrodes visualized by the mCherry expression (left panel). Right panel: extracellular voltage signal traces. **(D)** Time dependent visualization defined by category of the number of spikes per electrode for *BRAF*^*V600E*^, *BRAF*^*V600E*^/*p*Akt, *BRAF*^*V600E*^/*p*Akt/*Tp53*^*KO*^ and control slices incubated with aCSF. Time bin = 10 sec. Connecting lines are color-coded according to the category they belong to. **(E)** Representative traces of spontaneous firing from slices incubated with aCSF for 80 μsec. **(F)** Inter-model comparison of the mean of number of spikes per electrode from acute slices incubated with aCSF (*N* = 3 - 6; *n* = 7 - 21). **(G)** Inter-model comparison of the mean number of spikes per electrode of brain slices incubated with aCSF (low Mg^2+^, high K^+^); one-way ANOVA followed by Tukey′s multiple comparison test. **p < 0.01.

As brain slices are an isolated preparation lacking any sensory input, the levels of neuronal activity are much lower than *in vivo*. To probe how the distinct tumors entities responded to elevated activity we recorded spikes in brain slices bathed in a solution favoring network activity (aCSF low Mg^2+^, high K^+^). This maneuver could not increase activity in the tumor core regions of all three models (except for the control group) (**Fig. 5G**; *red* bars), suggesting that the central cytoarchitecture in the *BRAF*^*V600E*^ and *BRAF*^*V600E*^/*p*Akt/*Trp53*^*KO*^ neoplasms does not allow regular neuronal firing and that also in the *BRAF*^*V600E*^/*p*Akt tissue the dynamic range of neuronal activity is compromised compared to control (**Fig. 5G**; *red* bars). *BRAF*^*V600E*^ and *BRAF*^*V600E*^/*p*Akt/*Trp53*^*KO*^ tumors in the peri-tumoral area also did not respond to the solution exchange and thereby dampen activity infiltrating this region. Strikingly, we observed that the activity was substantially amplified in this transitional zone of *BRAF*^*V600E*^/*p*Akt mice and even exceeded the level of increase seen in control mice (**Fig. 5G**; *yellow* bars). Together with ongoing spontaneous activity in the core region, this suggests that the *BRAF*^*V600E*^/*p*Akt tumor model exhibits the strongest epileptogenic propensity.

## Discussion

Despite representing a seminal discovery, the presence of *BRAF*^*V600E*^ as prominent somatic mutation in developmental brain tumors demands further explanation with respect to (*a*) the large variety of affected glial and glioneuronal entities including PLNTY and GGs ^3,5^ and (*b*) differences in their biological behavior with a controversial role of *Trp53* ^14–17,39^. Our present data indicate that the heterogeneity and excitability of *BRAF*^*V600E*^-induced murine developmental brain tumors critically depends on the activity status of PI3K/mTOR pathway signaling and that *Trp53*-loss has fundamental impact on the acquisition of malignant features.

Of note, only *BRAF*^*V600E*^ on its own but not *p*Akt or *Trp53*-loss revealed the potential to induce neoplasia (**Table 1**). Thus, *BRAF*^*V600E*^ represents the primary oncogenic driver whereas *p*Akt and loss of *Trp53* confer specific pathogenetic and functional tumor features. Augmented PI3K/mTOR pathway signaling in concert with *BRAF*^*V600E*^ appears as precondition for the acquisition of *glioneuronal* features with frank dysmorphic neurons and neoplastic astroglia in emerging neoplasms thereby reflecting human GGs ^13^.

In contrast, *BRAF*^*V600E*^ alone introduced to neural precursors at E14 elicited MAP2-/Olig2-positive oligoid tumors that histologically reflect key differentiation aspects of PLNTY. In two large human series, glial features with prominent oligodendroglial characteristics as well as the presence of either *BRAF*^*V600E*^ or mutually exclusive fusion events involving FGFR2/FGFR3 were reported for PLNTY ^5,40^. The different molecular modifications converge in activating MAPK signaling. Of note, glio*neuronal* lesions have been demonstrated to arise from embryonal IUE-mediated transfer of GLAST-*BRAF*^*V600E*^ and Nestin-*BRAF*^*V600E*^, respectively ^34^ as well as in *BRAF*^*V637E*^-transgenic mice virally transfected with episomal Cre plasmid (**Table 1**) ^9^. Thus, the particular progenitor target subpopulation and the distinct developmental timepoint of somatic mutation may further increase the heterogeneity of *BRAF*^*V600E*^-induced tumors.

A further difference of the model approach in our present study is given by the use of truncated *BRAF*^*V600E*^ containing kinase domain. Truncated *BRAF*^*V600E*^ introduced by retroviral vector into neonatal Ntv mice under Nestin-promoter control induced tumors resembling pilocytic astrocytoma ^41^. Full length *BRAF*^*V600E*^ did not exert comparable effects potentially due to increased negative regulation of BRAF activity through possible phosphorylation of inhibiting residues on C-terminus domain in later progenitors pool available at birth or Hsp90 stabilizing binding in the full length BRAF^V600E^ protein ^42–45^.

The notion that retroviral transfer of *BRAF*^*V600E*^ in combination with an activated Akt variant into brains of newborn mice results in malignant tumors recapitulating features of (giant cell) glioblastoma multiforme (GBM) is as well compatible with the targeting of a postnatal progenitor population with glial fate and a different growth potential^46^. Intriguingly, our present data suggest that *BRAF*^*V600E*^/*p*Akt introduced to neural precursor populations at E14 generate benign neoplasms most closely resembling GG. Thus, the distinct progenitor population subjected to robust MAPK *and* PI3K/mTOR signaling is critical for both, neuropathological appearance and biological grade of the emerging tumors.

*BRAF*^*V600E*^/*p*Akt tumors require only a single additional genetic hit given by *Trp53* loss to acquire anaplasia. The malignancy of these tumors is also documented by their rapid growth and short survival of mice compared to other high grade neuroepithelial tumor variants in mouse models including invasive high-grade gliomas, medulloblastoma or adult glioma ^47–49^. Since *BRAF*^*V600E*^/pAkt/*Tp53*^*KO*^ neoplasms still encounter pronounced dysmorphic neurons, these tumors recapitulate clear anaplastic GG features, rather than characteristics of (epitheloid) GBM or anaplastic Xanthoastrocytoma. Multicolor labeling of tumor cells indicates that the *BRAF*^*V600E*^/pAkt/*Tp53*^*KO*^ neoplasms are multiclonal and derive from pluripotent neural precursors capable of giving rise to dysplastic neurons and neoplastic astroglia.

The transcriptional profile of the *BRAF*^*V600E*^/*p*Akt/*Trp53*^KO^ tumors strongly reflects their malignant biological behavior and invasive growth. Furthermore, pronounced transcript signatures relate to immune processes. Thus, augmented transcript patterns in *BRAF*^*V600E*^/*p*Akt/*Trp53*^KO^ tumors share major GO-term signatures with human malignant gliomas of *mesenchymal subtype* ^35^, which frequently harbor mutations of *Trp53* and *NF1*, the latter resulting in aberrant Ras/MAPK-*and* mTOR-pathway activation ^50^. Furthermore, there is reduced expression of transcripts related to synaptic structural homeostasis and transmission as well as potassium and calcium channels, neurotransmitter receptors and GABA-ergic inhibition despite the presence of a substantial dysmorphic neuronal components in *BRAF*^*V600E*^/*p*Akt/*Trp53*^KO^ neoplasms. Compromized inhibition related to reduced expression of GABA-homeostasis relevant molecules including GABA receptor and chloride potassium symporter transcripts has also been previously demonstrated in the microenvironment of gliomas ^36^. Thus, dysregulation of inhibitory circuits represents a common phenomenon in different brain tumors ^7^.

With respect to electrical activity, the present data suggest spike frequencies in the core region resembling activity in preexisting cortex as unique feature for the *BRAF*^*V600E*^/*p*Akt tumors. This might be unexpected as the cellular architecture is characterized by dysmorphic neurons and neoplastic astrocytes and thus differs from normal cortex. In this context, the findings by Goz and colleagues may be seminal demonstrating that *BRAF*^*V600E*^ expressed under GLAST-/Nestin promoters can confer a hyperexcitable phenotype to pyramidal neurons ^34^. We propose that this cellular hyperexcitability of *BRAF*^*V600E*^-positive neurons functionally compensates for the distorted network architecture to yield almost normal spike rates in the tumor core region. Spikes generated in this core tumor area will have a high propensity to propagate to neighboring cortical areas because the peri-tumoral region of *BRAF*^*V600E*^/*p*Akt tumors amplifies activity (**Fig. 5G**). As a consequence, the combination of maintained central neuronal activity with an enhanced relay to adjacent normal tissue will render this model most susceptible for the emergence of epileptiform activity.

In contrast, both 1- and 3-hit neoplasms lack significant intrinsic spike activity, which appears in line with the predominant glial differentiation of the former and a chaotic, dysfunctional cellular composition combined with reduced expression of transcripts instrumental for neuronal signaling in the latter.

Future studies will have to decipher the roles of secreted factors and synaptic inputs from the tumor environment present in diffuse gliomas ^37,38^ for the *BRAF*^*V600E*^-induced tumor spectrum. Our present data explain the molecular basis underlying heterogeneity of *BRAF*^*V600E*^-positive brain tumors and provides vistas for tailored therapy development.

## Acknowledgements

We thank Dr. D. Jones and Prof. J. LoTurco for providing plasmids.

