## Supplementary Data for "Heterogeneity and excitability of *BRAF*^*V600E*^-induced tumors is determined by PI3K/mTOR-signaling state and *Trp53*-loss"

### Supplementary Materials and Methods

**Experimental animals.** CD1 mice were obtained from Charles River (Strain code #022; RRID: IMSR_CRL:022) and B6.129P2-*Trp53*$^{tm1Brn}$/J mice were obtained from Jackson Laboratories (Stock No: #008462; RRID: IMSR_JAX:008462) and maintained under sterile conditions on a 12 h light cycle and fed *ad libitum*. To increase the IUE efficiency, B6.129P2-*Trp53*$^{tm1Brn}$/J mice were backcrossed with CD1 mice for three generations to generate *Trp53*$^{loxP/loxP}$ mice with a CD1 genetic background. For accelerated backcrossing, the genomes of the F1, F2 and F3 generations were sequenced, and mice with the lowest C57BL/6 single nucleotide polymorphism (SNP) score were used for further breeding. Since the CD1 background is predominant (estimated CD1 genetic content 85-90%), we further designate them as CD1-*Trp53*$^{loxP/loxP}$. In order to guarantee a reproducible genetic background for all experimental animals, we crossed female CD1-*Trp53*$^{loxP/loxP}$ mice and male C57BL/6-*Trp53*$^{loxP/loxP}$ mice to induce brain tumors. Mice without a severe clinical outcome were sacrificed at the age of 110 days, whereas mice affected with a severe phenotype after loosing 20% of body weight.

**Plasmids and cloning.** The binary piggyBac (PB) transposon system (Wellcome Trust Sanger Institute; Cambridge, UK) was used to integrate the transgenes into the genome of mice. A general donor plasmid (PB-CAG(CMV early enhancer/chicken β actin)-MCS) was generated by replacing the human UbC promoter from the PB-UbC-piggyBac-plasmid by the CAG promoter (kindly provided by Dr. J. LoTurco, University of Connecticut). In addition, a multiple cloning site (MCS) (5’-agcttagatctcccgggaccggt-3’) was added to the 3’ end of the promoter. From this general donor plasmid, we subsequently generated the donor plasmids PB-CAG-mCherry-lynAkt, PB-CAG-mCherry-BRAF$^{V600E}$ and PB-CAG-Cre-mCherry. For PB-CAG-mCherry-lynAkt, the *Akt* sequence from the myr-Akt1-pUSEamp (Addgene #17245; RRID: Addgene_17245) was amplified and subcloned into a plasmid containing mCherry-T2A (Addgene #72264; RRID: Addgene_72264). Next, a tyrosine-protein kinase Lyn sequence (gaccatggggtgtattaaatcgaaacggaaagacaacctt) was added to the 5' end of the *Akt* gene sequence to target Akt protein to the membrane. This gene will be further referred to as *p*Akt. The mCherry-*p*Akt cassette was then inserted into the MCS of PB-CAG-MCS by *HindIII* and *EcoRI* restriction sites (see primer sequences in **Supplementary Table S2**). For PB-CAG-mCherry-BRAF$^{V600E}$, the *BRAF$^{V600E}$* kinase domain (KD) gene sequence (kindly donated by Dr. David Jones, German Cancer Research Center, Heidelberg) was first subcloned into the plasmid containing mCherry-T2A (Addgene #72264; RRID: Addgene_72264) and subsequently inserted into PB-CAG-MCS using the same strategy as described above. In order to generate the donor plasmid PB-CAG-Cre-mCherry, the Cre sequence was amplified from AAV-GFP-Cre (Addgene #49056; RRID: Addgene_49056) and cloned into the MCS of PB-CAG-MCS-mCherry.

**Generation of Brainbow construct.** Brainbow is based on the Cre-*loxP* recombination system and insertion of multiples copies of the Brainbow allele into neural progenitors allows the target cell to express one color, which leads to the appearance of the daughter cells in the same color [18]. A plasmid containing pCAG-Brainbow3.0 was purchased from Addgene (#45176; RRID: Addgene_45176). Brainbow3.0 (encoding mOrange2, eGFP and mKate2 flanked by *loxP* sequences) was amplified and inserted into the MCS of PB-CAG-MCS by *BamHI* and *EcoRI* restriction sites (see primer sequences in **Supplementary Table S2**). We verify the newly cloned Brainbow construct *in vitro* and *in vivo* (**Supplementary Fig. S3B** and **C**),

**Generation of iRFP$^{713}$ construct.** iRFP$^{713}$ DNA sequence encoding the near-infrared fluorescent protein was amplified from piRFP (Addgene #31857; RRID: Addgene_31875) and digested by using *HindIII* and *AgeI* restriction enzymes. The resulting fragment was inserted into the MCS of PB-CAG-MCS, generating the donor plasmid PB-CAG-iRFP$^{713}$, further designed as CAG-iRFP$^{713}$ (see primer sequences in **Supplementary Table 2, Online Resource**).

**NS20Y cell transfection.** Mouse neuroblastoma derived NS20Y cell line was purchased from Merck (#08062517; RRID: CVCL_0469) and cultured in Dulbeco´s minimum essential medium (DMEM; Sigma) supplemented with 1% fetal bovine serum (FBS) and maintained at 37°C in a humidified incubator with 5% $CO_2$. NS20Y cells were transfected using Lipofectamine2000 (Invitrogen) one day after the cells were seeded. Medium was replaced to reduced serum medium (Opti-MEM; Life Technologies) 1 h prior cell transfection. DNA constructs (250 ng/well) and 1 µl of lipofectamine were diluted separately with 50 µl of Opti-MEM and incubated for 5 min at room temperature (RT). Both solutions were mixed and further incubated for 20 min at RT before its addition to the Opti-MEM medium containing the cells. After 24 h, the medium was replaced for DMEM containing 1% FBS.

**Western blot analysis.** Cell medium was removed after 48 h of cell transfection and phosphate-buffered saline (PBS) was added to wash the cells. Lysis buffer (150mM NaCl, 0.5% 50mM Tris-HCl, pH 7.4, 0.1% SDS, 1% sodium deoxycholate and 1% Nonidet P-40) with fresh added protease and phosphatase inhibitor cocktails (Millipore) was added to the cells and the cell homogenates were collected. Samples were incubated for 1 h at 4 °C, centrifugated at 10000 rpm for 10 min at 4 °C and mixed with Laemmli buffer prior denaturation at 95 °C for 5 min. Proteins were separated in 12% polyacrylamide-SDS gels and transferred in a nitrocellulose membrane. Blots were then blocked with 5% non-fat dry milk in 1% Tween diluted in Tris-buffered saline (TBS-T) for 1 h before the incubation with the primary antibody against pS6 and S6 for 2 h. Primary antibodies were diluted 1:1000 in 2% non-fat dry milk in TBS-T. After a washing step, membranes were incubated with an infrared-(IR)Dye-conjugated secondary antibody (LI-COR Bioscience) for 45 min. Proteins were visualized with the IR laser-based Odyssey Imaging System (LI-COR Bioscience). The antibodies used for western blotting shown in **Supplementary Table 4.**

**HEK293T cell transfection**. Human embryonic kidney (HEK 297 T) cell line was obtained from Agilent (Cat #240073; RRID: CVCL_6871) and continuously maintained in DMEM (Life Technologies), supplemented with 10% FBS and 1% penicillin/streptomycin, and incubated at 37°C in the $CO_2$ incubator. Transfection of HEK293 cells was performed by calcium phosphate ($CaPO_4$) method. Medium was replaced to 1 ml Iscove´s modified Dulbecco´s medium (IMDM;Life Technologies; supplemented with 5% FBS) per well. After mixing the DNA (500 ng/µl) with 50 µl 250 mM $CaCl_2$, 55 µl 2xHEBES phosphate-buffer (50 mM HEPES, 280 mM NaCl, 1.5 mM $Na_2PO_4$) was added to the DNA mixture while vortexing.

**Microscopy and imaging**. HE- and IHC-stained brain sections were scanned using a Mirax Scan digital microscope slide scanner (3DHistech) and the resulting images were visualized using the Case Viewer 2.3 Software (3DHistech). Epifluorescent images were collected with a confocal microscope (Nikon Eclipse Ti). Tumor slices were visualized using a staining with a secondary antibody conjugated with a fluorofore. Z-stack images were acquired at 1 µm intervals.

**Semiquantitative analysis of immunohistochemical sections.** Within the neoplastic region, microscopic areas of 9.4 $mm^2$ were photographed using a digital camera attached to a light microscope. Labeled (L) and non-labeled (NL) cells were quantified according to the DAB signal and judged as positive when the intensity of the staining was stronger than background. Cells in blood vessels or in their walls were excluded from the analysis [19]. Cell Counter Plugin of *Image J* software was used to quantify L and NL cells. The labeling index (LI) was quantified by dividing the L cells by the total number of cells (L+NL).

**KEGG pathway analysis and GO enrichment.** Kyoto Encyclopedia of genes and genomes (KEGG) pathway enrichment was performed using The Database for Annotation Visualization and Integrated Discovery (DAVID v 6.8). Pathways without any biological link to the genes were excluded from the analysis. To interpret the molecular function of the DE genes, Gene Ontology (GO) enrichment analysis was performed using the Gene ontology enrichment analysis and visualization tool (GOrilla) and FDR corrected p-values of 0.05 were considered significant.

**Histology and immunohistochemistry of human and mouse FFPE brain tissue**. For histological analysis, all tissue sections were stained with hematoxylin (Waldeck) and eosin (Roth; HE). For immunohistochemical (IHC) analysis, sections were deparaffinized prior unmasking with 10 mM citrate buffer (Sigma Aldrich, pH 9.0). Endogenous peroxidase activity was quenched with 1% $H_2O_2$ for 30 min. Slices were blocked in 5% normal goat serum (NGS; Thermo Fischer Scientific) in PBS for 30 min at 25 °C and incubated for 30 min at 4 °C with the primary antibodies. Samples were subsequently treated with either biotinylated anti-mouse or anti-rabbit (Vector Laboratories) antibodies, which were detected using a Vectastain kit (Vector Laboratories) and diaminobenzidine (DAB) staining (Sigma Aldrich). A counterstain was performed with hematoxylin incubation.

**Immunofluorescence stainings**. After mice decapitation, brains were removed and sectioned at 100um by using the vibratome (Thermo Fisher Scientific) and the resulting free-floating brain sections were incubated in 4% formaldehyde (Sigma Aldrich) overnight at 4 °C. After washing with PBS, slices were blocked with 0.1% Triton-X, 0.1% Tween-100 and 2% Bovine serum albumin (BSA) for 1 h at 25 °C prior to the overnight incubation with the primary antibodies. After washing in PBS, slices were incubated with the secondary antibody conjugated with an Alexa Fluor (Thermo Fisher Scientific) and DAPI (Sigma-Aldrich) for 2 h at 25 °C. After washing in PBS, free-floating brain slices were mounted using Mowiol (Sigma Aldrich). The antibodies used for immunohistochemistry are in listed in **Supplementary Table S3**.

**RNA extraction**. Tumor tissues and adjacent normal tissues from mouse brains were microdissected, frozen in liquid nitrogen and stored at -80 °C before RNA extraction. RNA was isolated using the RNeasy Micro Kit (Qiagen) following the manufacturer´s instructions and evaluated using a NanoDrop1000 spectrophotometer and a Qubit RNA HS Assay Kit on a Qubit 2.0 Fluorometer (Thermo Fisher Scientific).

**Bioinformatic analysis**. All bioinformatics analyses were performed on a linux system and R/Bioconductor packages Rsubread, edgeR and limma. After quality control and trimming (low quality tails, adapter and poly(A)-contamination) the raw sequencing data were mapped to the GRCm38/mm10 mouse and GrCh38/hg38 genome assemblies using the STAR aligner v.2.4.0.1. The read count matrix per gene was generated from the aligned sequencing data using the corresponding ENSEMBL transcript annotation files and featureCounts v.1.5.3. After filtering genes with low read counts and normalization, 17,511 genes were tested for differential expression. Linear models were fitted on the normalized expression data. Group-wise comparisons of the different tumor and control tissues were conducted. Raw p-values were adjusted for multiple testing using Benjamin-Hochberg. A FDR threshold of 5% was chosen to identify differentially expressed genes. Principal component analysis (PCA) and volcano plots were generated to visualize the differentially expressed genes of each comparison. Finally, unsupervised hierarchical clustering of the 100 most variable genes was performed to identify any gene expression patterns between the tumor and control tissue.

**Acute slice preparation.** Mice were decapitated shortly after anaesthetizing them with isoflurane. Brains were removed and glued on the vibratome tray in ice-cold carbogen-bubbled cutting solution containing (in mM) 87 NaCl, 2.5 KCl, 1.25 $NaH_2PO_4$, 7 $MgCl_2$, 0.5 $CaCl_2$, 25 $NaHCO_3$, 25 D-glucose and 75 sucrose, and further cut with the vibratome (Thermo Fisher Scientific). The resulting slices (300 µm) were maintained in the carbogen-bubbled cutting solution at 35 °C in an interphase chamber. After 30 min, the slices were transfered into an interphase chamber containing carbogen-bubbled artificial cerebrospinal fluid (aCSF) containing (in nM) 124 NaCl, 3 KCl, 1.25 $NaH_2PO_4$,

2 $MgCl_2$, 2 $CaCl_2$, 26 $NaHCO_3$ and 10 D-glucose, and incubated for at least 1 h before the recording.

**MEA recording and analysis.** Multielectrode array (MEA) plate consists of 64 electrodes with a diameter of 50 μm and arranged in an 8 x 8 grid (2.1 mm x 2.1 mm) (CytoView-MEA-6-well plate, Maestro Edge, Axion Biosystems). Brain slices were visualized with a fluorescence microscope (Leica M165 FC) in order to place the fluorescent tissue onto the electrodes. After the recording, an inverted microscope (Axio Observer 1A, Zeiss) was used to take pictures from position of the slices on the electrodes. AxIS software was used to export the number of spikes per electrode. Spikes were detected when the signal crossed the threshold of 6 x standard deviation of the baseline.

### Supplementary Figures and Tables

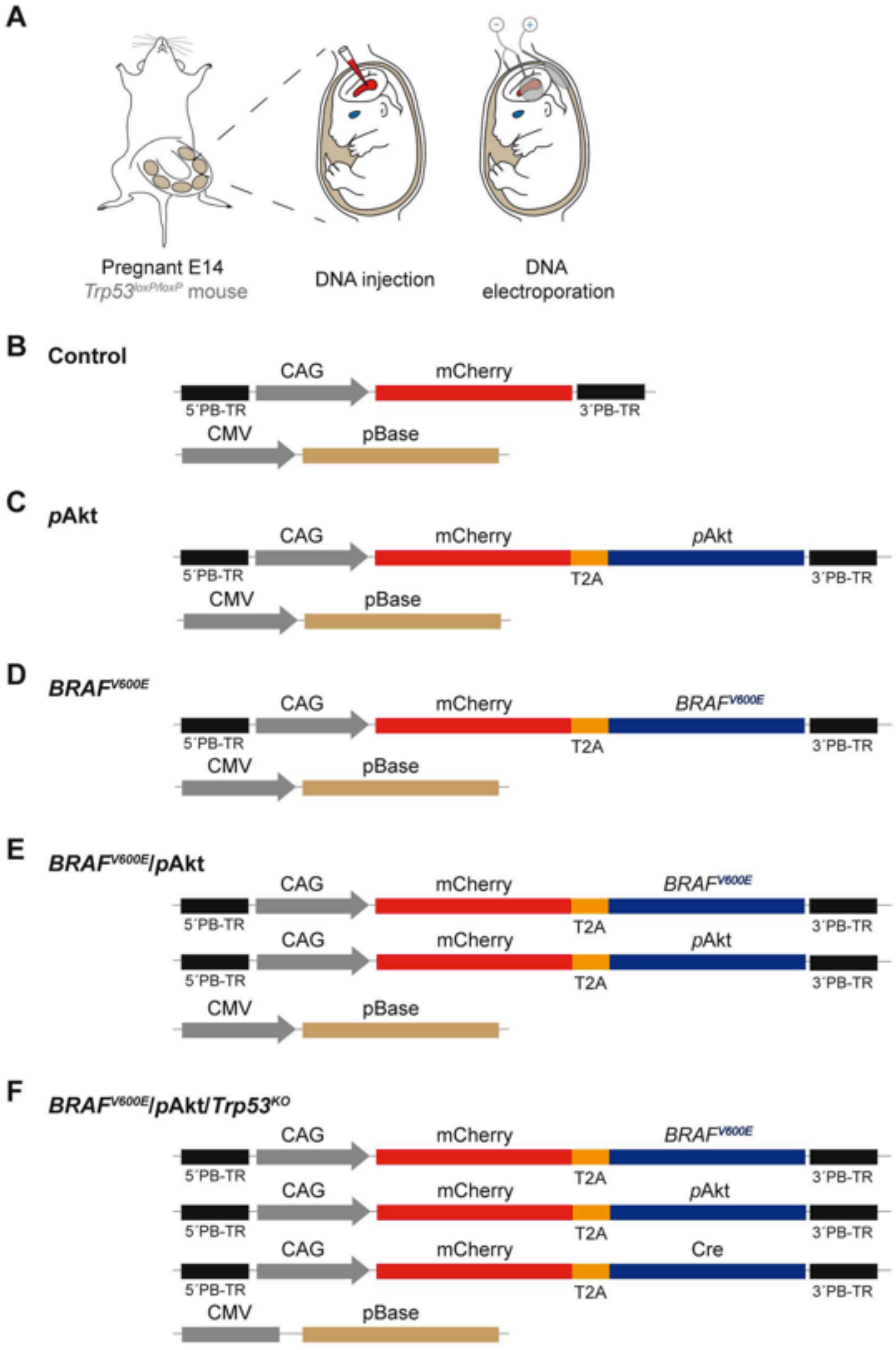

**Supplementary Fig. S1 Summary of the *in utero* electroporation (IUE) technique and schematic representation of the piggyBac plasmids used for IUE. Related to Figure 1-5. A,** Schematic drawing illustrating the IUE technique. Following uterus exposure at embryonic day 14 (E14), the DNA was injected directly into the third ventricle and 5 electrical pulses with an intensity of 40V and a length of 50 ms were applied. **B**-**F,** Schematic representation of the piggyBac donor and helper plasmids used for in IUE. In donor plasmids, piggyBac terminal repeats (PB-TR) flank the sequence subsequently integrated into the genome. The insertion of this sequence to the genome is possible due to the co-IUE of the helper plasmid CMV-pBase transposase. Note that the oncogenes are always co-expressed with the red fluorescent protein mCherry. The plasmid combinations used for IUE are the following: (**B**) PB-CAG-mCherry (Control), (c) PB-CAG-*p*Akt (*p*Akt) (**D**) PB-CAG-$BRAF^{V600E}$ ($BRAF^{V600E}$), (**E**) PB-CAG-$BRAF^{V600E}$ and –pAkt ($BRAF^{V600E}$/*p*Akt), (**F**) PB-CAG-$BRAF^{V600E}$, -*p*Akt and –Cre in $Trp53^{loxP/loxP}$ mice ($BRAF^{V600E}$/*p*Akt/$Tp53^{KO}$).

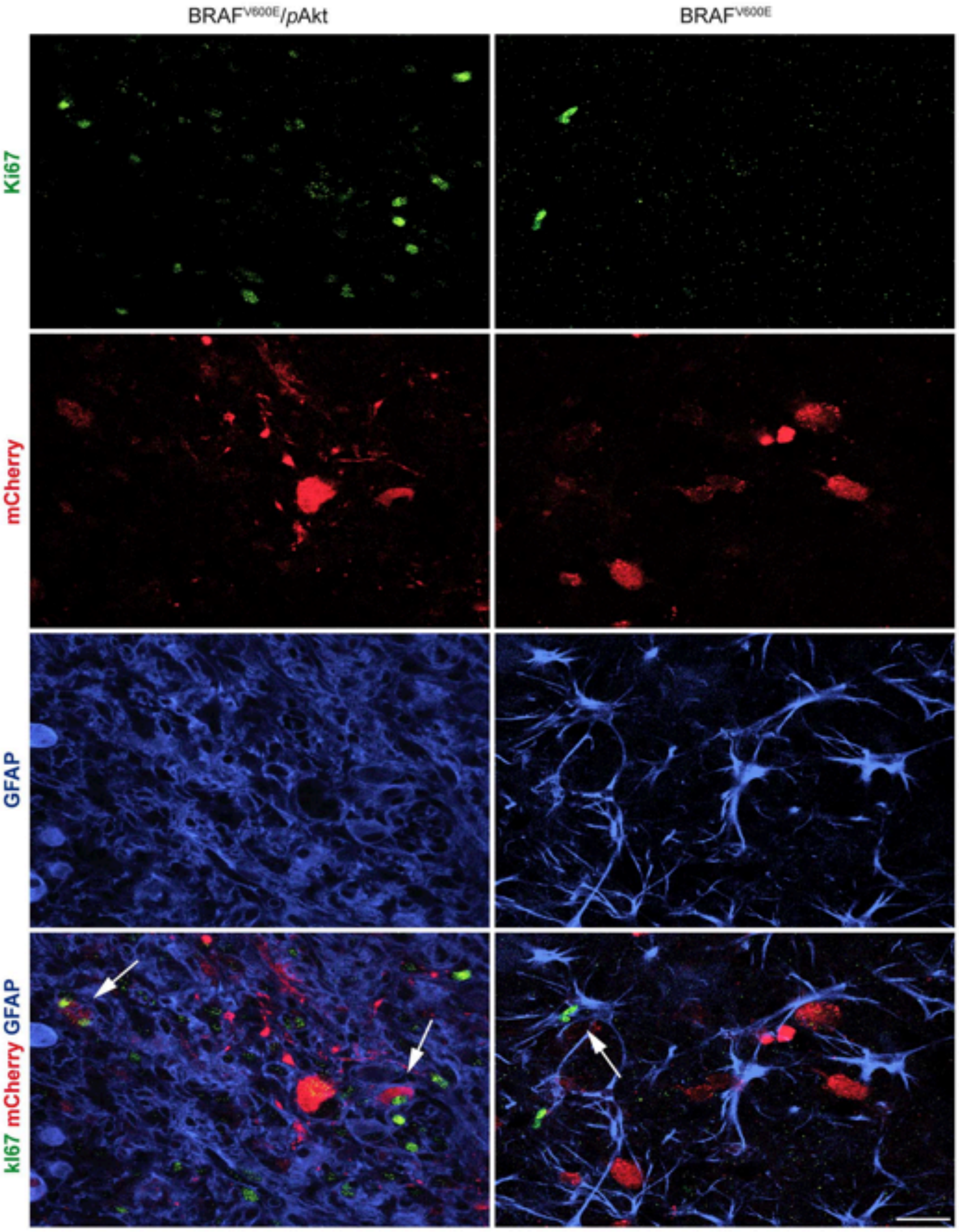


**Supplementary Fig. S2 Evaluation of the tumorigenic involvement of the astroglial component in *BRAF$^{V600E}$* and BRAF$^{V600E}$/*p*Akt tumors. Related to Figure 1.** BRAF$^{V600E}$/*p*Akt neoplasms but not BRAF$^{V600E}$ tumors contain a neoplastic glial component. Co-expression of the proliferation-associated antigen Ki67 and GFAP by astrocytes targeted by the IU-electroporated oncogenes (expressing mCherry) is observed only in $BRAF^{V600E}$/*p*Akt neoplasms but not in $BRAF^{V600E}$ tumors. Arrows point to a *reactive* astrocytic element with a delicate, isomorphic thus reactive process pattern in the $BRAF^{V600E}$ lesion versus more pleomorphic *neoplastic* astrocytic elements positive for mCherry in the $BRAF^{V600E}$/*p*Akt tumor. Scale bars, 25µm

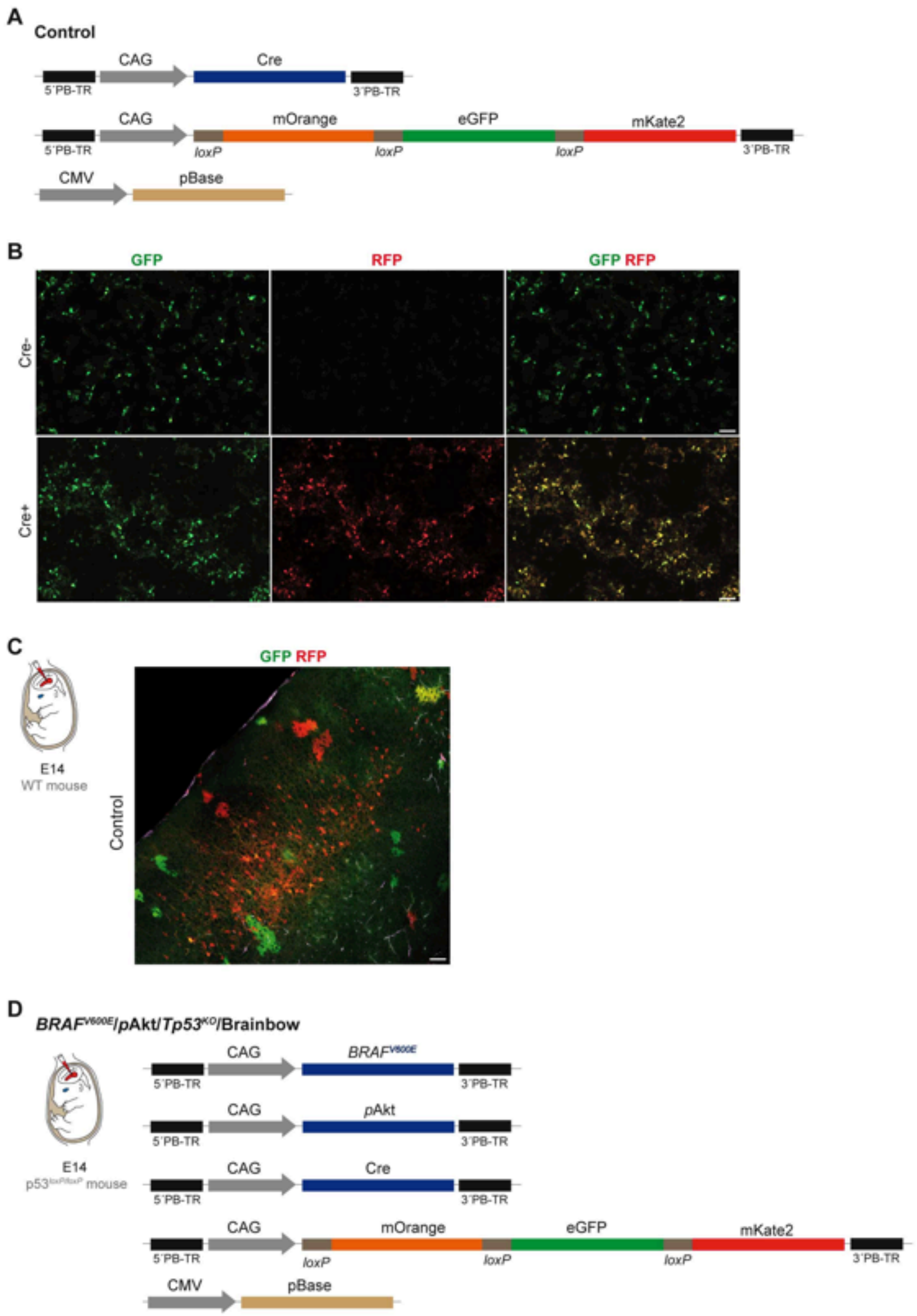
A
Control
CAG
Cre
5´PB-TR
3´PB-TR
CAG
mOrange
eGFP
mKate2
5´PB-TR
loxP
loxP
loxP
3´PB-TR
CMV
pBase
B
GFP
RFP
GFP RFP
Cre-
Cre+
C
GFP RFP
E14
WT mouse
Control
D
BRAF^V600E/pAkt/Tp53^KO/Brainbow
CAG
BRAF^V600E
5´PB-TR
3´PB-TR
CAG
pAkt
5´PB-TR
3´PB-TR
CAG
Cre
5´PB-TR
3´PB-TR
CAG
mOrange
eGFP
mKate2
5´PB-TR
loxP
loxP
loxP
3´PB-TR
CMV
pBase
E14
p53^loxP/loxP mouse

**Supplementary Fig. S3 Scheme of the combination of plasmids encoding Brainbow and oncogenes used to IUE at E14. Related to Figure 2. A,** Structure of the used donor plasmids PB-CAG-Cre and –Brainbow and the helper plasmid encoding the pBase transposase under the control of the CMV promoter. This combination is referred as control. PB-CAG-Brainbow encodes for three different fluorescent proteins (mOrange, eGFP and mKate), which are flanked by *loxP* sites. Cre transposase randomly recognizes and excises loxP sites leading to the target cell and its lineage to express a specific combination of fluorescent proteins. **B,** Micrographs showing HEK 293 cells transfected with the Brainbow construct with and without Cre transposase. Scale bar, 100 µm. **C,** Image showing a cortical area from a mouse (P40) IU-electroporated (E14) with the control plasmids (n = 5). PiggyBac transposition of Brainbow allows the targeting of neurons and glial cells with distinct fluorescent markers and the visualization of the clonally related cells, which share the same color combination. Scale bar, 100 µm. **D,** Structure of the piggyBac transposons and the helper plasmid pBase transposase used for IUE in $Trp53^{loxP/loxP}$ mice at E14 for the generation of neoplasms. The sequences flanked by the PB-TR are CAG-$BRAF^{V600E}$, -*p*Akt, -Cre and -Brainbow ($BRAF^{V600E}$/*p*Akt/$Trp53^{KO}$ + Brainbow). Note that the oncogenes are not co-expressed with a fluorescent tag when combined with Brainbow allele.

**Supplementary Table S1.** List of important differentially expressed genes in *BRAF*$^{V600E}$/pAkt/*Trp53*$^{KO}$ compared to control (ctrl).

***BRAF*$^{V600E}$/pAkt/*Trp53*$^{KO}$ vs ctrl**

| Gene symbol | logFC | FDR |
|---|---|---|
| *Scn1a* | -2.261202492 | 3.76E-05 |
| *Neurod1* | -2.882576707 | 0.001159647 |
| *Kcnab2* | -2.605519806 | 1.36E-08 |
| *Camk2b* | -2.090316674 | 2.84E-08 |
| *Syt13* | -2.833153877 | 3.05E-09 |
| *Lgals1* | 3.314488406 | 2.61E-09 |
| *Tnc* | 5.055604017 | 2.92E-08 |
| *Rest* | 2.760773242 | 5.21E-06 |
| *Mmp10* | 4.936537152 | 1.97E-06 |
| *Ccl22* | 2.188754314 | 0.003190229 |
| *Gfap* | 2.130149822 | 4.69E-05 |
| *Mki67* | 3.053196402 | 0.005726214 |

**Supplementary Table S2.** List of primer sequences used to clone the piggyBac donor plasmids.

| DNA sequence | Enzyme | Primer sequence 5´-3 |
|---|---|---|
| CAG-MCS_fw | *XhoI* | gcgctcgagtcattagttcatagcccatatatgg |
| CAG-MCS_rv | *BamHI* | gcgggatccaccggtcccgggagatctaagcttgccaaaatgatgag acagcacaa |
| myrAkt_fw | *BamHI* | gcgggatccatggggagcagcaagagcaa |
| myrAkt_rv | *EcoRI* | gcggaattcggctgtgccactggctgagt |
| mCherry-pAkt_fw | *HindIII* | gcgaagcttggaccaggaatggtgagcaagggcgagg |
| mCherry-pAkt_rv | *EcoRI* | gcggaattcggctgtgccactggctga |
| mCherry-BRAF$^{V600E}$_fw | *HindIII* | gcgaagcttggaccaggaatggtgagcaagggcgagg |
| mCherry-BRAF$^{V600E}$_rv | *EcoRI* | gcggaattcgtggacaggaaacgcaccat |
| mCherry_fw | *BamHI* | gcgggatccatggtgagcaagggcgaggagc |
| mCherry_rv | *EcoRI* | gcggaattcctagaagcccttgtacagctc |
| Cre_fw | *HindIII* | gcgaagcttatggccaatttactgaccgtac |
| Cre-(2A)_rv | *SmaI* | gcgcccgggggaagcggagagggcagaggaagtcttctaacatgc ggtgacgtggaggagaatcccggccctgatcgccatcttccagcagg c |
| Brainbow3.0_fw | *BamHI* | cgggaccggtggatcacaagtttgtacaaaaaagcaggc |
| Brainbow3.0_rv | *EcoRI* | ggcacagtcggaatttgagagacacaaaaaattccaacacactat |
| iRFP$^{713}$_fw | *HindIII* | tcattttggcaagcttatggctgaaggatccgtcg |
| iRFP$^{713}$_rv | *AgeI* | tcatggatccaccggttcactcttccatcacgccgatct |

**Supplementary Table S3.** List of the antibodies used for immunohistochemical and immunofluorescence stainings

| Antibody | Antigen | Dilution | Source | Identifier |
|---|---|---|---|---|
| Synapsin I | Synapsin I | 1:200 | Synaptic Systems | Cat #106011; RRID: AB_2619772 |
| GFAP | Glial Fibrillary Acidic Protein | 1:500 | Sigma Aldrich | Cat #G3893; RRID: AB_476889 |
| MAP2 | Microtubule Associated Protein 2 | 1:200 | Synaptic Systems | Cat #188002; RRID: AB_2138183 |
| Ki67 | Ki67 | 1:50 | Abcam | Cat #16667; RRID: AB_302459 |
| pS6 | Phospho-S6 Ribosomal Protein (Ser235/236) | 1:400 | Cell Signaling Technology | Cat #2211; RRID: 331679 |
| Olig2 | Olig2 | 1:20 | R&D Systems | Cat#AF2418; RRID: 2157554 |
| mCherry | mCherry | 1:300 | Abcam | Cat#167453; RRID: AB_2571870 |
| NeuN | Neuronal Nuclei | 1:200 | Merck | Cat #MAB377; RRID: AB_2298772 |

**Supplementary Table S4.** List of the antibodies used for western blot.

| Antibody | Antigen | Dilution | Source | Identifier |
|---|---|---|---|---|
| S6 | Ribosomal protein S6 | 1:1000 | Cell Signaling Technologies | Cat #2217; RRID: AB_331355 |
| pS6 | Phospho-Ribosomal protein S6 | 1:1000 | Cell Signaling Technologies | Cat #62016; RRID: AB_2799618 |
